# *In Situ* Unzipping of Long, High Molecular Weight DNA in Solid State Nanopores

**DOI:** 10.1101/2020.07.21.214445

**Authors:** R. Losakul, D. E. Tobar, A.M. Pfeffer, A. Gutierrez, R. Schipper, W. A. Jehle, H.W.Ch. Postma

## Abstract

Nanopores are an established paradigm in genome sequencing technology, with remarkable advances still being made today. All efforts continually address the challenges associated with rapid, accurate, high-throughput, and low cost detection, particularly with long-read length DNA. We report on the *in situ* melting and unzipping of *long, high molecular weight* DNA. At varying salt concentration, we directly compare the translocation conductance and speeds between SiN and graphene nanopores at sub-10 nm pore diameters. We observe the force-induced unzipping of dsDNA at higher salt concentrations than previously reported in literature. We observe free running translocation without secondary structures of ssDNA that is an order of magnitude longer than reported before. We hypothesize that the frayed single strands at the molecule’s end get captured with a higher likelihood than both ends together. In understanding this phenomena for long-read lengths, we continue to address the challenges revolving around future generations of sequencing technology.

**Statement of Significance:** Genome sequencing is an advancing field with applications in clinical diagnostics. However, the challenges of providing accurate identification of longer DNA molecules at low cost are still developing. While detection of long DNA molecules is established, the identification of its individual nucleotides presents its own set of challenges. By separating the hydrogen bonds between the two strands, individual nucleotides are made directly measurable. However, identification is hindered from the formation of secondary structures, where the single-stranded DNA sticks to itself. Previous studies only included short DNA molecules. We report in situ force-induced unzipping and translocation of long DNA without secondary structures almost an order of magnitude longer than reported before. Our findings present new experimental conditions and insights that progress the field towards high accuracy sequencing of individual long molecules.

## I. INTRODUCTION

Nanopore biosensors are a rapidly advancing technology, investigating the unknowns of human genetics and genome biology. With potential applications in clinical diagnostics or personalized medicine, nanopores utilize electrophoretically driven DNA molecules through a trans-membrane channel. This technique provides immediate and direct measurements of nucleic acids by their electrical current [1]. Biological nanopores have shown success in single-molecule detection, in both academic and commercial settings, using trans-membrane proteins like alpha-hemolysin [2], with automated ratcheting for precision [3], and more recently, *de novo* sequencing for longer read lengths (100 − 882 kb) [4]. Additional studies propose platforms integrating force spectroscopy over an array of nanopores accompanied with avidin-biotin anchors, removing the need for polymerase chain reaction (PCR) assays [5–7]. However, due to the structural instability and natural lifetimes of proteins, commercial scalability and long-term storage are limited.

Solid state nanopores are a more robust alternative and offer mechanical, thermal, and chemical stability with pore size customization (for a review, see Dekker [8]). Additionally, novel fabrication techniques like focused ion-beam sculpting [9], TEM ablation lithography [10–12], and dielectric breakdown [13] have been created and successfully replicated. Previous studies with ultra thin materials like silicon nitride (SiN) and graphene have demonstrated these advantages in conjunction to dsDNA translocations. Similar to protein pores, solidstate nanopores also act as a transmembrane channel between two chambers of electrolyte solution. When electrophoretically driving DNA, an ionic current flows in response to an applied transmembrane electric field. Every translocated nucleotide perturbing the electric field generates a distinct blockage current and is immediately recorded.

In order to progress towards single-molecule *sequencing*, accurate detection of individual nucleotides is paramount. Achieved through DNA denaturation, the separation of hydrogen bonds between dsDNA backbones, single-stranded DNA (ssDNA) is required to detect and read nucleotides. Previous studies of DNA denaturation have shown dependence on melting temperature when treated with varying temperature, chemical solvents, and salt concentrations [14, 15]. With initial theory on the mechanical stability, entropy, and heat capacity changes, additional studies report a force-induced unzipping of dsDNA during translocation [16, 17]. This was experimentally demonstrated and confirmed in protein pores with PCR analysis [18] and subsequently demonstrated in synthetic pores. Notably with SiN membranes at sub-2 nm pore diameters, sequencing is indicated along with unzipping kinetics. These studies report an Arrhenius barrier dependence on duplex length [19] and distinctive three-level current blockades [20]. At slightly larger pore diameters (4 − 10 nm), single-stranded translocations was also observed with unfolded and unraveled configurations [21, 22]. However, these reports where conducted with short duplex DNA comprised with unstructured homo-polymers, hairpin molecules, or hybrid structures with single-stranded overhangs. In order to address the challenges of rapid, high accuracy, high throughput, and low cost detection of *long* read-length DNA, further understanding of long DNA interactions is required.

In this paper, we present *in situ* melting and unzipping of long, high molecular weight lambda-DNA in both SiN and graphene nanopores. Directly comparing between SiN and graphene membranes, we report the changes in conductance values, translocation duration and speeds at varying salt concentrations with sub-10 nm pore diameters. We observe low salt translocation of ssDNA with minimal secondary structures. More importantly, we also observe mechanical unzipping of dsDNA, at higher salt concentrations than expected from melting temperatures reported in literature [14]. We hypothesize that at low *n*_KCl_, the dsDNA begins to fray [23, 24]. Due to the reduced persistence length *L*_*P*_, these frayed ends have a higher likelihood of capture in the nanopore over the common mode double strand. Furthermore, under a strong electric field, *in situ* unzipping would occur as DNA molecules translocate. In understanding this phenomena, we can continue to address the challenges in solid-state devices and progress towards new generations of sequencing technology.

## II. MATERIALS & METHODS

### A. Materials

Solid SiN membranes were acquired from Norcada Inc (NX5004Z-60O). They consist of a 5 *×* 5 mm^2^ Si frame with thickness of 200 µm, a (20 − 30 µm)^2^ membrane with thickness 12 nm, and a 60 nm thick underlayer of SiO_2_. SiN membranes with a micropore for graphene studies were acquired from Norcada Inc (custom order). They either had a 5 *×* 5 mm^2^ or 3.5 *×* 5 mm^2^ Si frame with 200 µm thickness, a 50 µm membrane with thickness of 200 nm or 50 nm, respectively, and a 60 nm thick underlayer of SiO and a 5 nm overlayer of SiO for improved adhesion to graphene. Micropore diameters ranged from 0.5 − 2 µm.

### B. Device Fabrication

**Graphene samples** (Fig. 1) were made from mechanically exfoliated 5 − 10 mm size natural graphite (NGS Naturgraphit GmbH) flakes with Blue Nitto tape (Nitto Denko, SPV 224LB-PE). Graphite flakes were deposited on 300 nm SiO_2_/500 µm Si wafers that were previously cleaned in separate ultrasonic baths of Acetone (PHARMCO-AAPER), IPA (99%, PHARMCO-AAPER), de-ionized Water, and dipped in a 6:1 Buffered Oxide Etch (HF, JT.Baker 1178-03). Graphene flakes were identified with an optical microscope (Meiji MT7530 BF/DF), with thicknesses determined by optical contrast using a calibration curve obtained with Atomic Force Microscopy (Pacific Nanotech Dual Scan).

**FIG. 1.**
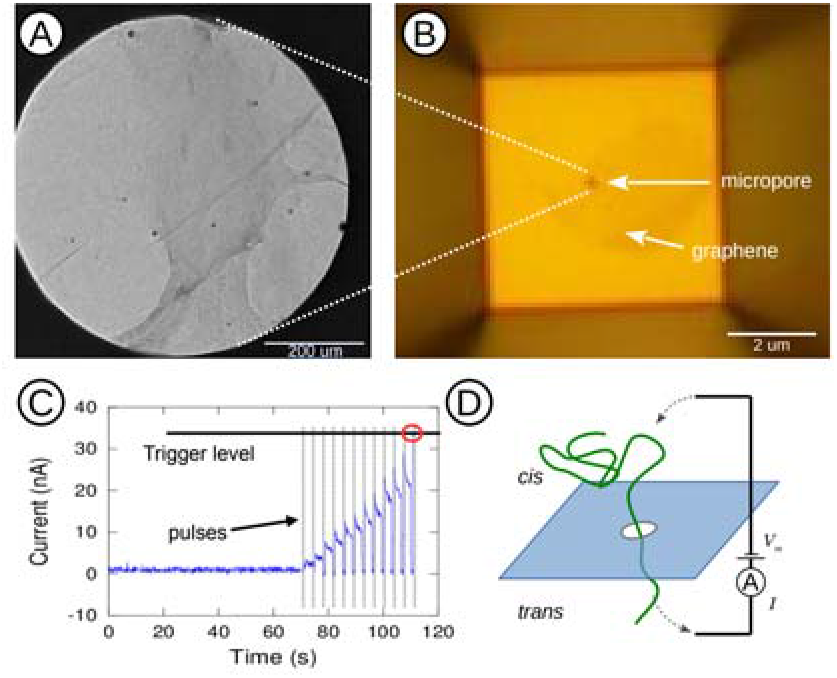
Fabrication of nanopores and DNA translocation procedure. A) Transmission Electron Micrograph of suspended graphene membrane over micropore. B) Optical image of graphene membrane over micropore. C) Nanopore fabrication procedure. For graphene nanopore, a pulsing procedure was used, which was terminated at reaching a predetermined trigger level. For SiN membranes, a constant voltage procedure was used (not shown). D) A transmembrane voltage *V*_*m*_ is applied and the ion current *I* (dashed) is measured as DNA (green) translocates through a nanopore from the *cis* to *trans* side.

We used the wedge transfer technique developed by Schneider *et al.* [25] to transfer and deposit individual graphene flakes onto micropores. We covered individual graphene flakes in a droplet of Cellulose Acetate Butyrate (CAB, Sigma-Aldrich 419036-250G) in Ethyl Acetate (EtAc, Sigma Aldrich 319902-1L) mixture, with its location manually marked. The graphene flake and CAB transfer polymer were wedged off from its Si/SiO_2_ substrate in de-ionized water with a freshly aspirated surface, before transferred onto a hydrophilic micropore treated with RF oxygen plasma for three minutes (Harrick Plasma PDC-32G, 18 W). Under an optical microscope with a modified stage, the graphene flake and CAB was manually positioned over the micropore before residual water completely dried. After ensuring the graphene was placed correctly (Fig. 1b), the micropore with graphene under CAB was placed on a hotplate at 75 °C for 30 min to promote secure adhesion. The CAB was dissolved in two separate EtAc baths for 30 s. The remaining graphene flake was deposited onto the micropore by annealing at 400 °C for 30 min. Suspended graphene samples were imaged in a Transmission Electron Microscope (FEI Titan S/TEM) (Fig. 1a).

**SiN samples** were acquired commercially and cleaned with Piranha Etch and treated with RF oxygen plasma.

A custom microfluidic flow cell was immersed in DI water with its channels pre-flushed to remove residual air. Assembled devices were immersed in ethanol (EtOH, PHARMCO-AAPER, 111000200) to remove trapped air inside the micropore pyramid. The pre-wetted device was transferred and mounted in the flow cell under water. Once removed from water, the mounted device was secured and the outside of the cell was dried thoroughly before both channels were flushed with 2 M KCl buffer solution for initial *I*(*V*) measurements.

### C. Nanopore Fabrication

Once membranes were confirmed to be insulating, nanopores were fabricated in both SiN and graphene membranes using dielectric breakdown[13, 26]. **Graphene pores** were formed by rapid pulses of increasing duration Δ*t* and height Δ*V*. Depending on the desired pore size, we simultaneously observed whether *R* exceeded 10 − 20 MΩ at 0.2 V. Initial pulses were Δ*t* = 1 µs long and Δ*V* = 3 V high. Then Δ*V* was slowly increased up to 10 V. If no initial pore was formed, Δ*V* was reduced and Δ*t* was increased. Nanopores formed at 6 − 10 V at 1 µs − 1 ms. Upon initial pore formation, pulses at 3 − 4 V of similar duration as initial formation were used to adjust the nanopore diameter *d* (Fig. 1d).

**SiN nanopores** were made by slowly ramping a *constant bias V* until a desired trigger current level *I* was reached. The pore was enlarged by resetting *V* and ramping up again, and the trigger level was reached at decreasing *V* levels. Initial trigger currents were *I* = 30 − 80 nA and initial formation occurred at *V* = 4 − 6 V. Post breakdown enlarging caused triggering to occur below ∼ 1 V.

An initial value of the nanopore diameter *d* was determined from the open pore conductance *G* using the bare pore conductance *G*_0_ and access resistance *R*_*a*_ [27, 28]

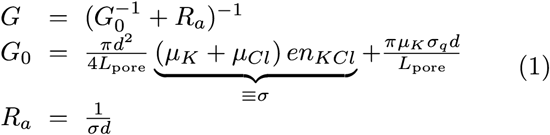

Note that *σ*_*q*_ is surface charge density while *n*_KCl_ conductivity is *σ*. The pore diameter *d* and depth *L*_pore_ were determined self consistently using *G*(*n*_KCl_ = 2 M) and Δ*G*(*n*_KCl_ = 2 M) [29].

### D. Buffers and DNA

2 M KCl buffer solution was made with 2 M KCl, 10 mM Tris HCl, and 1 mM EDTA. Additionally, Lambda DNA (N3011L, New England Biolabs) and KCl buffers with an initial pH ∼ 5. When changing *n*_KCl_ from 2 M to a lower value, we track *G vs. t* and wait until it saturates at a value consistent with Eq. 1. We do not see a contribution from *σ*_*q*_ in these experiments, which we comment on below. Experiments were performed at room temperature.

### E. Data Acquisition & Analysis

The *cis*-channel of the flow cell was flushed with lambda DNA solutions at selected KCl concentrations. The *trans*-channel flushed with selected KCl concentration only. A trans-membrane voltage *V*_*m*_ was applied to electrophoretically drive the DNA through the nanopore. For SiN at all *n*_KCl_ values *V*_*m*_ ≤ 100 mV, while *V*_*m*_ ≤ 200 mV for graphene nanopores. The corresponding ion current change was recorded (Fig. 2) with a data acquisition board (National Instruments PCI-6251) and custom acquisition software at 0.1 − 1.25 MS*/*s, with a Fourier filter to remove stray interference.

**FIG. 2.**
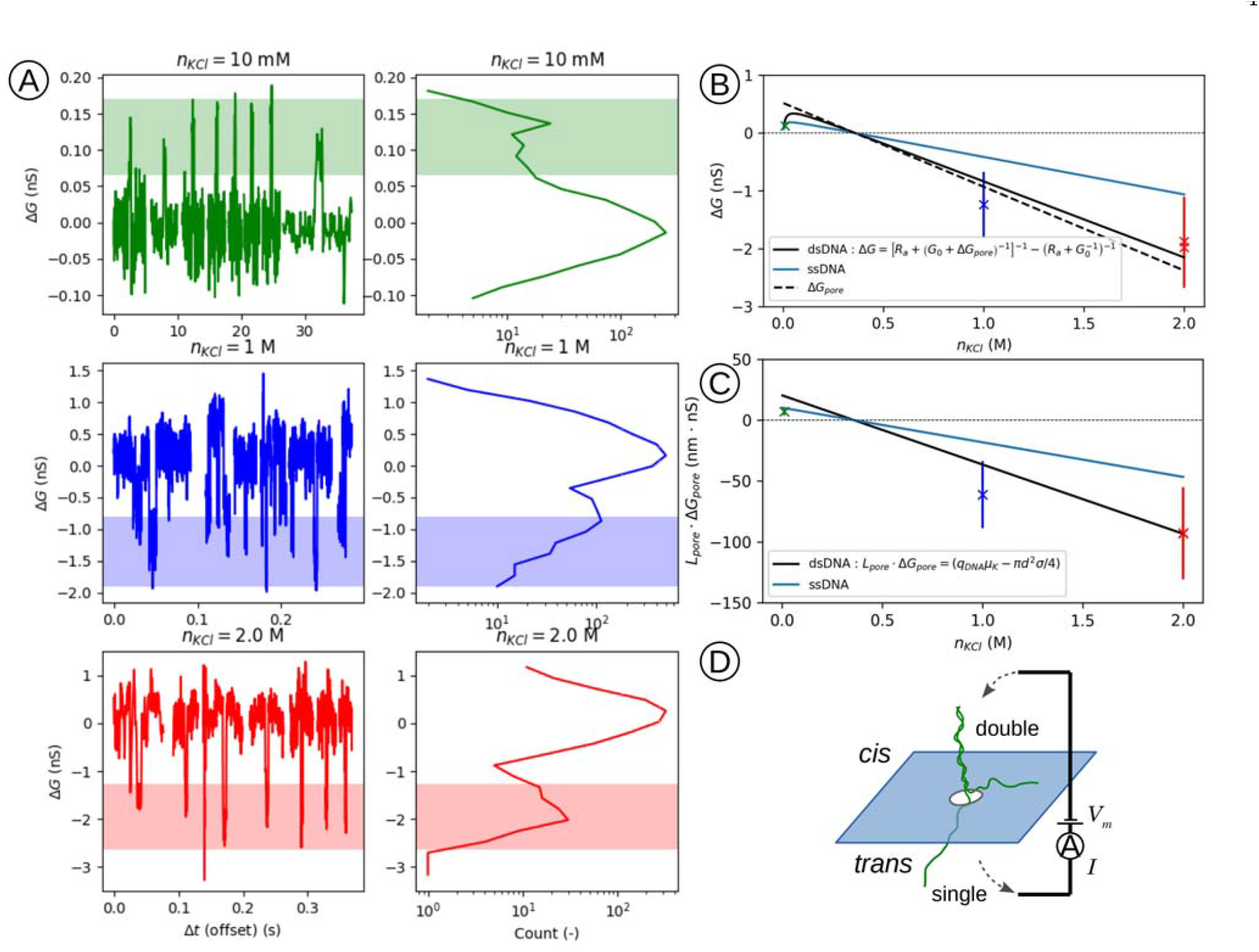
Unzipping of high-molecular-weight DNA through SiN nanopores. A) Left: Subset of Δ*G* for events at *n*_*KCl*_ = 0.010 (green), 1 (blue), and 2 M (red). Right: Histogram of all event current traces for corresponding conditions in left panel with secondary peak width (shaded). B) Expected mean of Δ*G* values of ssDNA (blue), dsDNA (black), and Eq. 2 (dashed) *vs. n*_KCl_. C) Expected mean of *L*_pore_ Δ*G*_pore_ for ssDNA (blue) and dsDNA (black) *vs. n*_KCl_. The 2 M (red) and 1 M (blue) align with dsDNA and the 10 mM (green) aligns with ssDNA. D) Diagram of dsDNA (green) unzipping, as it moves through a nanopore in a membrane (blue) from the *cis* to *trans* side driven by a transmembrane potential *V*_*m*_ while monitoring the current *I*.

To filter out low-current signal events in the presence of a high 1*/f* -noise background, common in nanopore systems [30, 31], we detected events as follows. The signal is leveled by subtracting a 100-point convolution average. The standard deviation is recorded for every 100-point section of the leveled signal, *i. e.* the background *I*_*bg*_, with the median standard deviation *σ*_*m*_ of that set recorded as well. Possible events were identified as exceeding *I*_*bg*_ by a multiple of *σ*_*m*_, without yielding false positives: 3 to 5 *σ*_*m*_. The signal was then convolved with a 2-point average, before the procedure repeated. Subsequently, events were fitted with the expected rectangular shape of height Δ*I* and duration *τ* of translocation waveforms, deduplicated, and reduced based on the *χ*^2^ values and signal to noise ratio Δ*I/σ*_*m*_.

This manner of event detection is particularly useful for detecting low-signal events against a high 1*/f* background. The convolution filter preserves the squareness of the event waveform so it is not distorted when large parts of the spectrum are rejected. Furthermore, the ratio of the shortest timebase *τ*_*s*_ (reduced timebase due to convolution) to longest timebase *τ*_*l*_ (100 point baseline subtraction) gives a constant bandwidth ratio that returns constant noise RMS values 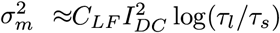 when 1*/f* noise dominates. This facilitates a numerical threshold criterion that does not depend on the part of the spectrum under consideration. For graphene nanopores we find *C*_*LF*_ ≈ 10^−5^ to 10^−3^ with no clear *n*_KCl_ dependence, while for SiN nanopores we find *C*_*LF*_ ≈ 10^−4^ to 10^−3^ at *n*_KCl_ = 2.0 M with a monotonic increase of a factor ≈ 10 down to *n*_KCl_ = 10 mM.

## III. OBSERVATIONS

We observe a series of short changes in the ion conductance Δ*G* across the nanopore (Fig. 2) every time a DNA molecule translocates through the pore [8]. To ensure these fluctuations are well separated from the baseline current *I*_*bg*_ and display a distinctive peak in the ion current histogram, we limit the passage of folded molecules by fabricating relatively small nanopores, *d* = 4 *…* 7 nm. At high *n*_KCl_, we normally find Δ*G <* 0 because it is dominated by the exclusion of ions from the nanopores.

Conversely at low *n*_KCl_, Δ*G >* 0, since counterion current from the DNA backbone dominates [27]. Both effects are described by

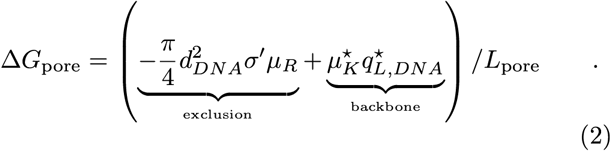

Due to the charge *q*^*\**^ and cross-section *πd*^2^*/*4 of dsDNA being halved during denaturation or unzipping, its conductance is also halved, 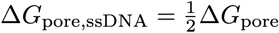. Over a range of *n*_KCl_ concentrations, the expected conductance values of dsDNA (black) and ssDNA (blue) are shown in Fig. 2 and are directly compared to our observations.

We observe two major deviations from Eq. 2 for SiN and graphene nanopores. The first deviation occurs between expected values between *n*_KCl_ and Δ*G*. With pore diameters between *d* = 5 − 7 nm, SiN nanopores observed at lower *n*_KCl_ had Δ*G* values significantly smaller than Eq. 2 predicts. For graphene nanopores with diameters between *d* = 4 − 7 nm, we observe an overall trend of Δ*G >* 0 at high *n*_KCl_ and Δ*G <* 0 at low *n*_KCl_. However, we also observe at *n*_KCl_ ∼ 220 mM, Δ*G <* 0 in contrast to the prediction of Δ*G >* 0 from Eq. 2 (Fig. 3).

**FIG. 3.**
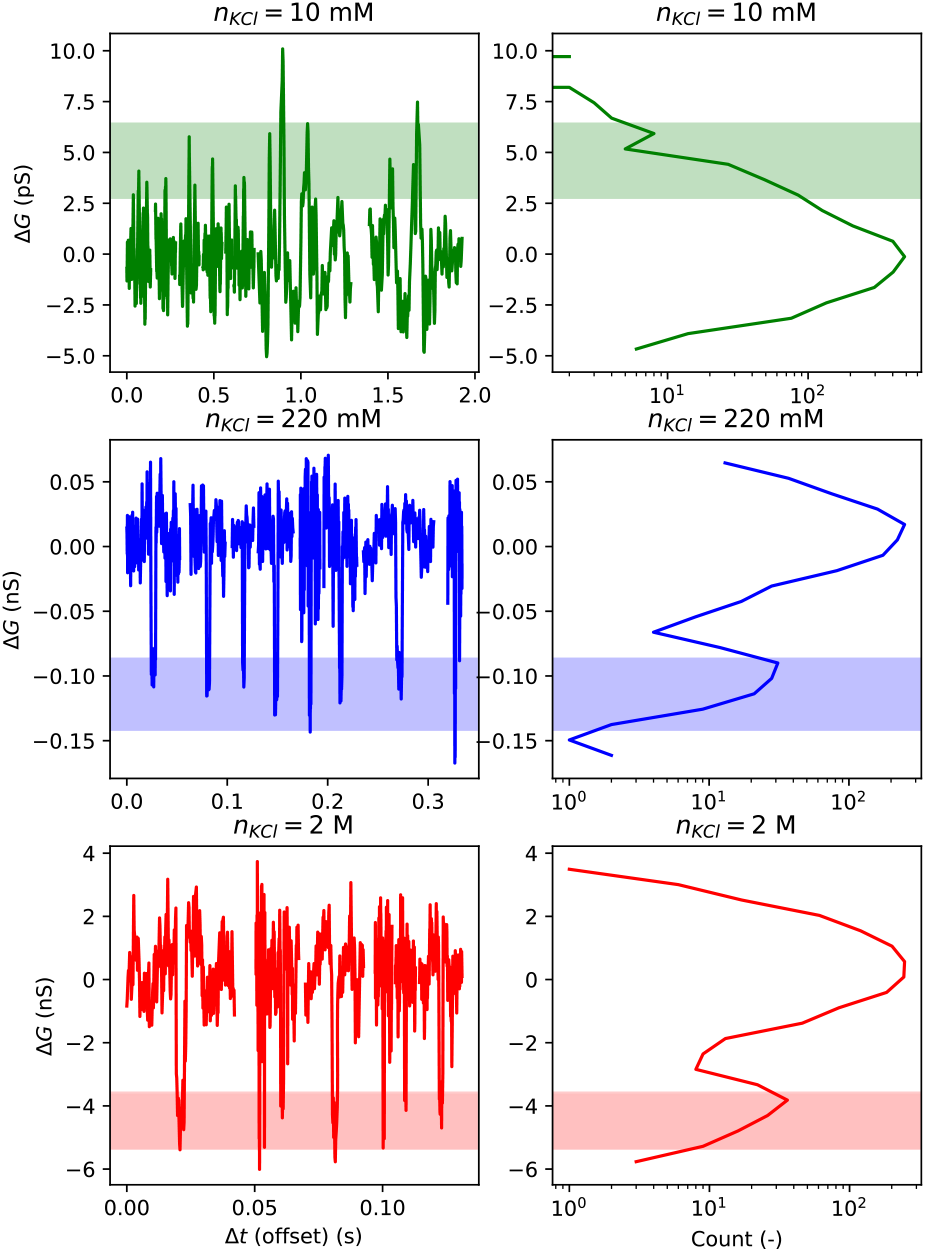
Unzipping of high-molecular-weight DNA through graphene nanopores. Left: Subset of Δ*G* for events at *n*_*KCl*_ = 0.010 (green), 1 (blue), and 2 M (red). Right: Histogram of all event current traces for corresponding conditions in left panel with secondary peak width (shaded).

The second deviation occurs with reduced mobility *µ*. While we designed our graphene nanopores to deliberately be treated *without* a passivation layer to inhibit ssDNA binding due to pi-stacking interactions [32], at *n*_KCl_ = 10 mM we see translocation events with Δ*G >* 0 and much reduced *µ* (Fig. 3, 4). Simultaneously, we see a slow decrease in background current *I*_*bg*_ and therefore an increase of background resistance *R* in a linear manner at a rate of d*R/*d*t* ≈ 10 %*/* min. We do not see an exponential dependence within ∼ 5 min. This increase is only visible at forward bias, *V*_*m*_ *>* 0, not reverse bias. When applying reverse bias pulses with duration and height typical for pore enlargement, the conductance does not recover. In contrast at higher *n*_KCl_, the background resistance is stable and there is an upper limit to the *R* drift, *i. e.* |d*R/*d*t* | ≲ 0.5 %. The increase of *R* at low *n*_KCl_ ultimately limits the duration of the experiment and lifetime of graphene devices, and we do not observe this with SiN nanopores.

**FIG. 4.**
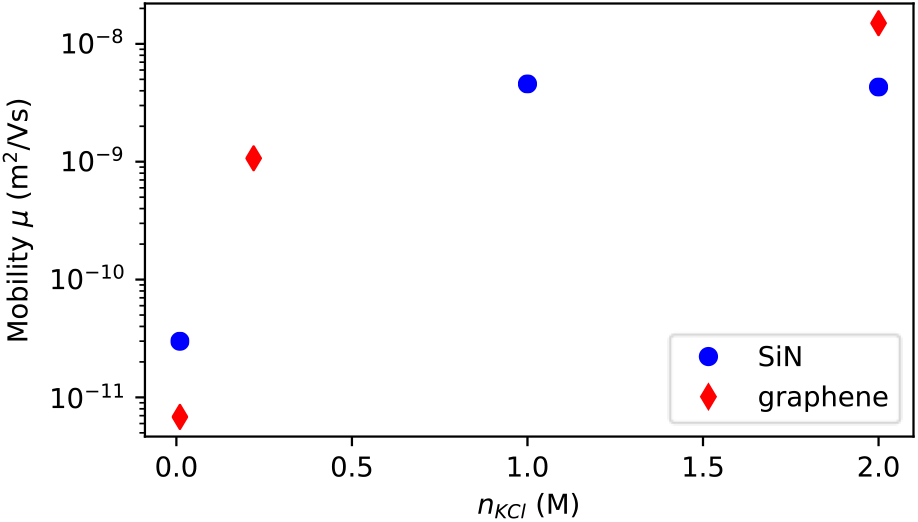
Mobility of DNA driven through graphene (red diamonds) and SiN nanopores (blue circles) as a function of *n*_KCl_.

For both SiN and graphene nanopores, we studied the mobility *µ* of DNA driven through the nanopore. Mobility was determined as *µ* = *v/E*, where *v* is the translocation speed and *E* the effective field in the nanopore (Fig. 4). We used *E* = *V*_eff_ */L*_pore_, where *V*_eff_ is the voltage drop across *G*_0_ + Δ*G*_pore_ in a series 1*/*(*G*_0_ + Δ*G*_pore_)+*R*_*a*_ network that divides down the applied voltage *V*_*m*_. Typical field strengths were 2 MV m^−1^ for SiN and 20 MV m^−1^ for graphene. At high ionic strengths, we find typical *µ* for DNA in both graphene and SiN nanopores [33]. However, at 10 mM, *µ* is reduced by a factor ∼ 100, for both SiN and graphene nanopores.

## IV. DISCUSSION

### A. SiN Nanopores

#### The role of R_a_

The values for Δ*G* at low *n*_KCl_ are much lower than Eq. 2 describes (Fig. 2). We argue that in a low *n*_KCl_ regime, the effect of access resistance *R*_*a*_ must be taken into account. When the pore is open, *R*_*a*_ and 1*/G*_0_ constitute a voltage divider, whereas when DNA is inside the pore, *R*_*a*_ and 1*/*(*G*_0_ + Δ*G*_pore_) constitutes one. If *d* ≪ *L*_pore_ at high *n*_KCl_, most of the voltage drops across the pore and Δ*G* follows Eq. 2. However, at low *n*_KCl_the geometric dependence of Δ*G*_pore_ changes and the condition *d* ≪ *L*_pore_ no longer guarantees *R*_*a*_ ≪ (*G*_0_ + Δ*G*_pore_)^−1^. Consequently, the quantity actually measured is

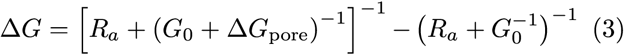

(Fig. 2b). We extract Δ*G*_pore_ using the inverse of Eq. 3, causing the ssDNA and dsDNA curves at low *n*_KCl_ to separate (Fig. 2c) and observe a distinct grouping of Δ*G*_pore_ on the ssDNA curves.

#### The role of *σ*_q_

We do not see a contribution of surface charge *σ*_*q*_ to the open pore conductance *G* of our nanopores. While this is expected for graphene, for SiN in our experiments *σ*_*q*_ ∼ 0 since we operated close to the charge neutrality point pH ∼ 4.1 [34]. Alternatively, we could have non-zero *σ*_*q*_ if *n*_KCl_ is different than expected. We use the settling of *G* to an expected value according to *n*_KCl_ as an indicator that we reached the right concentration. It can be argued that due to residual *n*_KCl_, a higher *n*_KCl_ is actually reached. However, that also *increases G*. Another argument could incorporate *σ*_*q*_ by assuming *n*_KCl_ is much lower than introduced. However, in our experience, exchanging high *n*_KCl_ for lower *n*_KCl_ only appears to lead to slightly *higher n*_KCl_ than desired, if fresh solvent is not used enough and/or we do not wait long enough. Pore shrinkage combined with a higher effective *n*_KCl_ together could allow for a finite *σ*_*q*_ that would explain our open pore conductance values. However, we confirm that SiN pores usually do not shrink by going back to the starting *n*_KCl_ and finding the same open pore conductance. [We note that if *n*_KCl_ were higher, it would also significantly reduce the unzipping capability]. A final alternative interpretation is that we have a source of contamination on the SiN membranes, despite our exhaustive cleaning the membranes with organic solvents, oxygen plasma, and Piranha. We calibrate the pore geometry as described and find *L*_pore_ = 20 − 40 nm larger than the bare membrane. If this is due to an accumulation of material that does not contribute to *σ*_*q*_ that would be consistent with our observations.

#### Halving of ΔG_pore_

As Δ*G*_pore_ is on the ssDNA curve (Fig. 2C), we suspect that we are translocating *single* strands despite starting with high-MW *double* strands. Under physiological conditions, the two strands of dsDNA bind together because the hydrogen bonds and stacking energies are stronger than the effective electrostatic repulsion of the two negatively charged backbones. This repulsion is overcome because the ca<u>tions screen th</u>e backbone with a Debye length 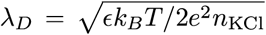, where *E* is the permittivity, *k*_*B*_ is Boltzmann’s constant, *T* is the absolute temperature, and *e* is the electron charge. At room temperature *λ*_*D*_(*n*_KCl_ = 1 M) = 0.3 nm while *λ*_*D*_(*n*_KCl_ = 10 mM) = 3.0 nm, which exceeds the distance between the backbones [35]. This effect favors the separation of the dual strands and lowers the melting temperature *T*_*m*_. However, *T*_*m*_ for high-MW dsDNA at *n*_KCl_= 10 mM is usually around 70 °C [14] and depends on sequence [36].

We hypothesize that we are unzipping the dsDNA due to the large *E* field at the pore. While the unzipping or force-induced melting of dsDNA is a well studied subject, the nanopores used previously are usually smaller than the dsDNA size (*<* 2 nm) and very short DNA (∼ 50 b) strands are typically employed [5, 6, 18–20, 37, 38]. When ∼ 5 kb strands are used, secondary structures cause large blockage signatures and a slowdown of the translocation process [22]. Here, we appear to drive *high-molecular-weight* ssDNA through a nanopore *without secondary structures*. The energetics of the unzipping process are usually analyzed using a Kramers-type 1D escape problem [5, 39]. An applied force *f* acts over a range Δ*x* in overcoming the barrier, such that

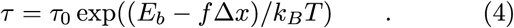

Using a hydrogen bond range of Δ*x* ≈ 0.6 nm [37], a field strength of *E* ≈ 8.3 MV m^−1^, an effective charge of 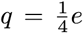 per base [40], we find *f* Δ*x* = 0.05 *k*_*B*_*T*. Once unzipping has started under physiological conditions, it takes ∼ 0.1 *k*_*B*_*T* [41] to break each additional pair, well within range of the available energy. Under this scenario, the capture rate could be limited by the initial energy barrier to get the unzipping going, afterwards the process is downhill.

#### Unzipping selection mechanism

We are using nanopores that are in principle large enough to accommodate both strands simultaneously. In fact, we use the dsDNA signal to calibrate *d* and *L*_pore_ *in situ*. Without a selection mechanism, dsDNA is as likely to translocate as unzipped ssDNA. How is it then possible that we appear to translocate unzipped single stranded molecules? First, since we see few, if any, folded and/or doublestrand signals, we assume the dominant capture mode is end-capture. In addition to small *d*, folded capture is also limited by the much larger persistence length *L*_*p*_ at low ionic strength [42–45]. Second, since we are close to *T*_*m*_, denaturing bubbles [46] are expected to cause a rapid time dependent fraying of the ends of the molecule [23, 24].

We put forward two hypotheses. First, the individual frayed strands at the molecule’s end are expected to fluctuate more than the common mode double strand fluctuations due to the much reduced *L*_*p*_ of ss over dsDNA, even at low ionic strength. If this constitutes an increase in attempt frequency for capture, this would favor unzipping. Second, if the pore is charged and that charge is poorly screened, it could renormalize *d* thus favoring passage of single over double strands. However, we do not see a role of *σ*_*q*_ for SiN pores and do not believe that surface charge of the pore plays a role in unzipping. We posit that the preferential trapping of the frayed single strands is the selection mechanism responsible for unzipping over double-strand translocation. A compounding mechanism could be envisioned that works similarly to geometric selection, *i. e.* denaturation bubbles are larger than fully hybridized dsDNA. Due to that size being close to *d*, dsDNA could be prevented from entering the pore. While the bubble size is most likely limited to the hydrogen bond length of 0.6 nm for few-base bubbles, it may be larger at the frayed end of dsDNA. Experiments at larger *d* could elucidate the role of this effect.

#### Grouping of translocation events

It is possibly expected that once a single strand has translocated, its complement would be captured quickly afterwards, which could cause a grouping in inter-translocation times. The two strands would translocate quickly after one another, followed by translocation of another molecule at a longer time. We do not see such a grouping which may be due to the large scatter in inter-event times.

### B. Graphene Nanopores

#### ΔG for graphene nanopores

Δ*G* does not appear to follow Eq. 2, especially at *n*_KCl_ = 220 mM where the sign of Δ*G* is opposite from that expected (Fig. 3). We argue that the particular geometry of our graphene nanopores have an effect on Δ*G*. Specifically, we have a significantly lower aspect ratio *α* = *L*_pore_*/d* in our graphene nanopores than in SiN nanopores. This causes *R*_*a*_ to dominate the nanopore resistance resulting in *G* ∝ *d*, instead of *G* ∝ *d*^2^ [12, 28, 47].

Continuum modeling shows that the current density *j*_*i*_ is carried mostly at the perimeter of the nanopore. Moreover, non-uniform *j* can also be expected due to the local dipole moments at the graphene nanopore edge. The crossover *n*_KCl_^⋆^ deduced from Eq. 2 assumes uniform *j*_*i*_ inside the nanopore. Since *j*_*i*_ is not uniform, but carried at the nanopore perimeter, we must compare the counter-ion current density *j*_*c*_ with *j*_*i*_ and integrate over the DNA cross-section. For small nanopores, such as the ones studied in this paper, a significant part of the DNA’s cross section is blocking the higher *j*_*i*_ at the pore edge. This would push 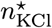 lower and could explain why we observe Δ*G <* 0. For larger nanopores and *α*, if the point of translocation is approximately in the center of the nanopore, the blocked *j*_*i*_ is *lower* than the uniform *j*_*i*_ and push 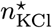 higher. However, a thorough experimental and theoretical evaluation of this effect has not been reported before and is beyond the scope of this report.

#### Selection mechanism for grapheme

Specifically for unzipping at graphene pores, the pi bonding could cause one of the strands to be tied to the graphene, freeing up the other strand for translocation. However, we see similar *µ* at low *n*_KCl_ with graphene, which does not carry a major surface charge save for some dipoles at the nanopore edge, *e. g.* carboxyl groups.

#### I_bg_ reduction at low n_KCl_ in grapheme

As the reduction does not appear in SiN pores at low *n*_KCl_ and only at forward bias with graphene pores, the decrease is consistent with a very slow binding of DNA material close to the pore. As the binding does not appear reversible by voltage pulsing, pi-bonding of free bases in denaturation bubbles [46] to the hexagonal graphene lattice is a logical candidate [48]. Close to the nanopore, we expect *E* to be stronger in graphene pores owing to the much smaller *L*_pore_, even when factoring in the larger role of *R*_*a*_. Furthermore, if unzipping occurs at SiN due to *E*, it is reasonable to expect it does so for graphene nanopores as well.

### C. Both SiN and Graphene Nanopores

#### Single nanopores

The breakdown method of making nanopores requires nucleation of a defect, followed by a slow opening of the pore through the same defect [13]. It is possible that we have more than a single defect and rather than opening a single defect, we are opening two or more defects. However, if we allow for a parallel current path and use that to solve for the pore geometry, we do not get self-consistent results. Therefore, we conclude we only measure single nanopores.

#### Mobility reduction

This may be expected due to the unzipping introducing a rate limit. We can quantify the rate with Eq. 4 but the attempt frequency is difficult to determine independently for the complicated nanoelectro-mechanical landscape in the vicinity of the pore and is subject of further studies. The mobility reduction can also be due to DNA adhering to the substrate and could explain the lower *µ* for graphene than SiN. Indeed, the ∼ 10 fold reduction for graphene at 220 mM could be due to DNA adhesion, as well. However, since the largest share of reduction in *µ* is similar for both types of pores, we hypothesize the unzipping itself is primarily responsible for lowering *µ*, as has been observed in space-constricted unzipping in smaller nanopores [49].

## V. CONCLUSIONS

We present evidence of unzipping high-MW DNA *in situ* due to the strong field at a nanopore entrance and translocating single strands without secondary structure, resulting in a ∼ 100-fold reduction in *µ*. Despite the pores being large enough to accommodate both strands, we preferentially capture single strands at the frayed ends of the DNA. We hypothesize that single end capture mechanism is due to an increased attempt frequency for capture for single over double strands owing to the much shorter persistence length.

For follow-up studies, we plan to explore the phase diagram of the unzipping process. First, raising the cell temperature to *T*_*m*_ should yield unambiguous melted DNA and therefore a clear Δ*G at the same n*_KCl_ that we can compare to. Second, the trade-off between *E* and proximity to *T*_*m*_ represents a 2D phase diagram in which more points can be chosen to determine the phase boundary. Third, adjusting the pH value to alter the surface charge of the SiN surface and allow an independent inspection of the role of *σ*_*q*_. Fourth, further exploring the unzipping speed scale with energy barrier once the bias is factored in, as a function of proximity to *T*_*m*_. Fifth, distinguishing event signals between blunt end and single-stranded overhang translocations. Finally, determining the role of larger nanopore diameters in unzipping and single-stranded translocation.

## VI. AUTHOR CONTRIBUTIONS

Conceptualization, RL, DT, HP; Methodology, RL, HP; Software, HP; Validation, AG; Formal Analysis, RL, RS, HP; Investigation, RL, DT, AP, AG, WJ, HP; Resources, RL, DT, AP, AG, WJ, HP; Data Curation, RL, RS, HP; Writing-Original Draft, RL, DT, HP; Writing-Review & Editing, RL, HP; Visualization, HP; Supervision, HP; Project Administration, HP; Funding Acquisition, HP.

## VII. ACKNOWLEDGEMENTS

The research reported here was supported by the National Human Genome Research Institute of the National Institutes of Health under Award Number R21HG10056. We are grateful for the experimental assistance in TEM imaging by William A. Hubbard and Jared J. Lodico from the Regan group at UCLA and discussions with Aleksei Aksimentiev, David Hoogerheide, Vincent Tabard-Cossa, Meni Wanunu, and Michael Zwolak.

